# Hypercomplex properties of *Drosophila* object detecting neurons

**DOI:** 10.1101/081398

**Authors:** Mehmet F. Keleş, Mark A. Frye

## Abstract

Many animals rely on vision to detect objects such as conspecifics, predators, and prey. Hypercomplex cells of the feline cortex and small target motion detectors of the dragonfly and hoverfly optic lobe demonstrate robust tuning for small objects with weak or no response to elongated edges or movement of the visual panorama [1–4]. However, the relationship between anatomical, molecular, and functional properties of object detection circuitry is not understood. Here, we characterize a previously identified lobula columnar neuron (LC11) in *Drosophila* [5]. By imaging calcium dynamics with two-photon excitation microscopy we show that LC11 responds to the non-directional motion of a small object darker than the background, with little or no responses to static flicker, elongated bars, or panoramic gratings. LC11 dendrites reside at the boundary between GABA-ergic and cholinergic layers of the lobula, each dendrite spans enough columns to sample 75-degrees of visual space, yet the functional receptive field is only 20-degrees wide, and shows robust responses to an object spanning less than one 5-degree facet of the compound eye. The dendrites of neighboring LC11s encode object motion retinotopically, but the axon terminals fuse into a glomerular structure in the central brain where retinotopy is lost. Blocking inhibitory ionic currents abolishes small object sensitivity and facilitates responses to elongated bars and gratings. Our results reveal high acuity small object motion detection in the *Drosophila* optic lobe.

## RESULTS AND DISCUSSION

Flies readily orient toward large moving objects such as elongated vertical bars or edges representing landscape features, and this behavior is mediated by interactions between motion vision and motion-independent object detection processes [6–10]. However, flies are able to perform object-directed behaviors when directionally selective columnar motion detectors (T4 and T5) supplying the third optic ganglion, the lobula plate, are silenced [7,8,11], suggesting the existence of object detection circuitry that acts independently from the motion vision pathway [12]. Recent studies have revealed many of the cells, circuits and computations for motion vision from photoreceptors to the lobula plate [13–19], yet the functional role of the neighboring neuropile, the lobula, which includes 80% of all neurons in the lobula complex [20], remains enigmatic in *Drosophila*.

Recent neuroanatomical and computational work has shown that the lobula contains more than 16 types of visual projection neurons (VPNs) that are columnar in the lobula (LC) and project to distinct synapse-rich output domains (optic glomeruli) in the ventrolateral protocerebrum (VLPR) [5,21]. Neurons downstream of LCs which interconnect multiple optic glomeruli in the central brain, have been shown to be sensitive to small objects [22], raising the possibility that some LCs themselves may be tuned to small objects. Optophysiological and electrophysiological methods have demonstrated that several LCs are broadly sensitive to visual features such as edges or bars [23,24], yet no study to date has explored LCs for the detection of small two-dimensional solid objects.

We visually-screened the publicly-available Janelia Gal4 lines [25] and identified a driver which, according to previous expression data [5], labels LC11 (Figure 1A). Analyzing single cell clones [26], we demonstrate that the R22H02-Gal4 driver labels ~51 LC11s (±4, n=5) and each LC11 neuron has dendritic arborizations in layers 2,3,4 and 5 of the lobula (Figure 1A,B, Figure S1A, A. Nern personal comms.) [5]. The dendrites cover 14-15 lobula columns in the dorso-ventral axis and 6-8 columns in the antero-posterior axis (Figure 1C and D). Assuming that each lobula column maps to a single ommatidium and taking into account that LC11 dendrites form an ellipse in the lobula (Figure 1E), we estimated the total span of a single LC11 to be 65 to 85 ommatidia thereby comprising a ‘multi-pixel’ columnar neuron. Considering the dendritic span (~15x8 columns), the number of LC11 cells (~50), and the array of total retinal ommatidia (~28×28), the dendritic arbors of individual LC11s very likely overlap one another extensively.

**Figure 1.**
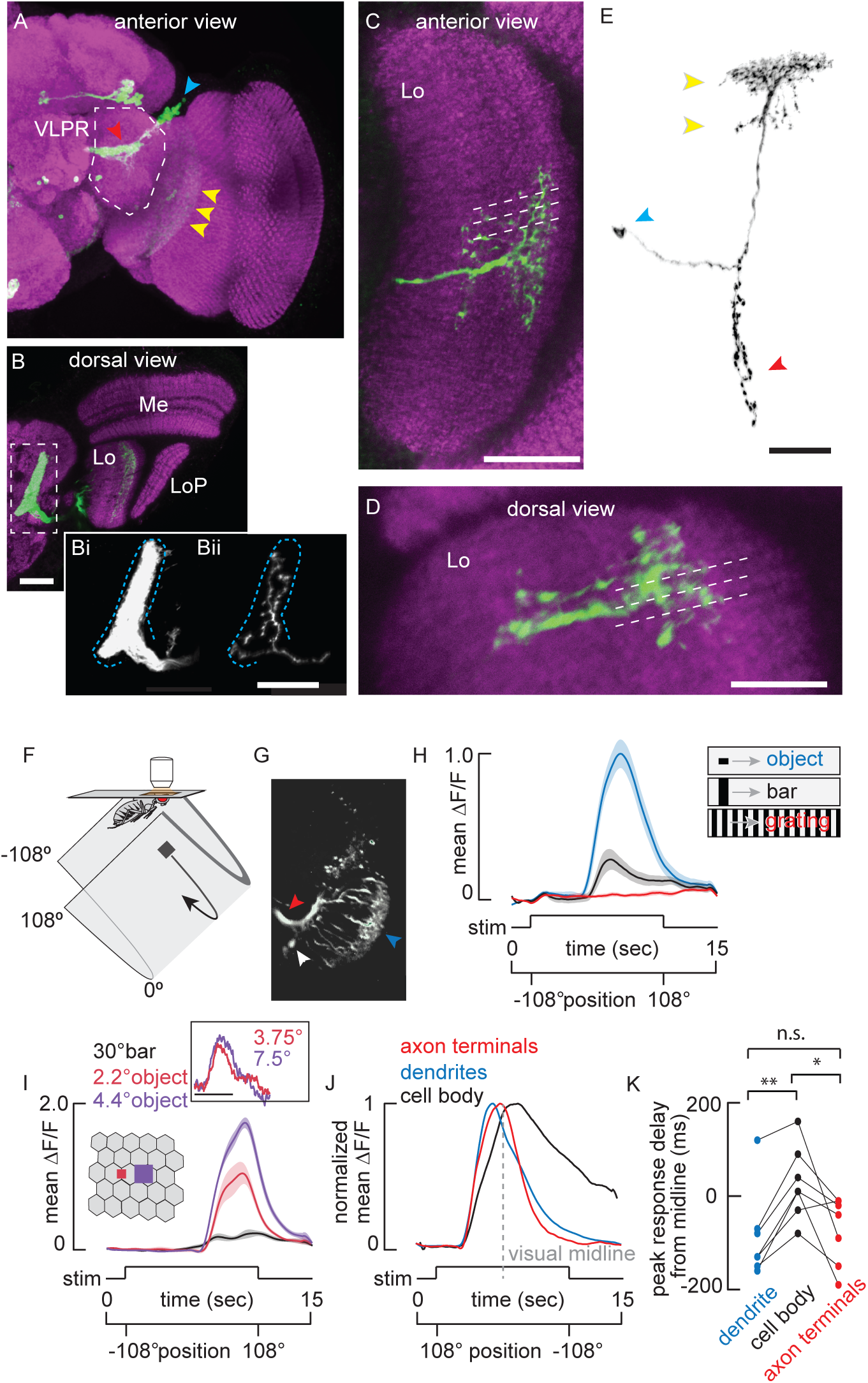
Anatomy of Lobula Columnar 11. A) Anterior view of a brain from a fly expressing membrane tethered GFP under the control of R22H02-Gal4 and labeled with anti-GFP (green) and anti-nc82 (magenta). The maximum intensity projection of a confocal stack was used to create the image. Dashed lines indicate the approximate boundary of the ventrolateral protocerebrum. Arrowheads indicate cellular compartments; blue-cell bodies, red-terminals, yellow-dendrites. B) Dorsally mounted view of R22H02-Gal4>UAS-mCD8::GFP flies. Dashed rectangle indicates the unique foot shaped LC11 glomerulus. Me: Medulla, Lo: Lobula, LoP: Lobula Plate. Comparison of the labeling of ~50 LC11s innervating the glomerulus (Bi) vs. stochastic labeling of a single LC11 (Bii). Single LC11 shows the full glomerular innervation with no evidence of retinotopic organization. Blue dashed line indicates the glomerular boundary. C) Anterior view of a stochastic labeling of a single LC11. Dashed lines indicate individual columns within the lobula. The dendritic arbor of a single LC11 covers about 14-15 lobula retinotopic columns spanning this axis. D) Dorsal view of a single LC11. Dendritic arbors span 6-7 columns spanning this axis. E) Morphology of a single LC11. Yellow arrows indicate the bistratified dendritic morphology of LC11 within lobula. Blue and red arrowheads indicate cell body and terminals respectively. All scale bars are 25 um. F) Schematic of the 2-photon imaging setup. The fly’s head is fixed and the surrounding LED arena covers 216° in azimuth and 70° in elevation. G) Image of LC11s expressing GCaMP6m under two-photon microscopy. Arrowheads indicate dendrites (blue), glomerular projections (red) and cell bodies (white). H) Mean GCaMP6m (±S.E.M. shading) signal of LC11 glomerulus in response to the movement of a 30°by 8.8° object (blue), a 30° by 70° bar (black) and a wide-field grating (red, n = 7 flies). I) Mean GCaMP6m (±S.E.M. shading) signal of the LC11 glomerulus in response to the movement of a 30° by 70° bar (black), 2.2° square object (red) and 4.4° square object (purple) (n = 6 flies). Inset: From tethered flies, normalized mean steering responses to single pixel impulsive displacement of objects, sizes indicated (scale bar: t=0-100 ms, n = 10 flies). J) Normalized mean ΔF/F of cell body (black), dendrites (blue) and axon terminals (red) responses to the movement of a small object as in C. Visual midline indicated with a dashed grey line. n = 7 flies. K) Comparison of the response delay between the dendrites (blue), terminal (red), and cell body (black) (*p<0.05, **p<0.01, paired t-test, n = 7 flies).

Previous work showed that all LC11 presynaptic sites are confined to the cognate glomerulus in the posterior VLP (PVLP) [5,21]. By contrast, we observe presynaptic sites not only in the glomerulus but also within layer 5 innervations of LC11 (Figure S1A). Whereas the synaptotagmin labeling was weaker in the lobula compared to the PVLP (data not shown), it did not overlap with a dendritic marker (Figure S1B). We can exclude the possibility that the synaptotagmin labeling in this layer was due to an additional cell in our Gal4 line because this cell does not innervate layer 5 (Figure S1B).

By contrast to the retinotopic organization of the columnar dendrites in the lobula, labeling a single LC11 neuron reveals spatially-distributed presynaptic terminals ramifying throughout the glomerulus formed by LC11 terminals (Figure 1B i-ii). This convergent organization suggests a loss of retinotopy at the glomerulus, consistent with findings in other LCs [21]. We investigated the significance of this convergence through a series of physiological experiments to elucidate the functional properties of LC11.

We utilized GCaMP6m under two-photon imaging (Figure 1F) to record LC11 dendritic, axonal and cell body responses (Figure 1G) to a small object that may represent another fly nearby or a larger animal moving at a distance, a vertical bar that may represent a landscape feature such as a tree, and a rotating wide-field grating representing optic flow generated during self-motion. We observed large responses to the small object within the axon terminals of LC11 (Figure 1H). The response diminished markedly for the vertical bar, and there was no response to the wide-field grating (Figure 1H).

Flies can behaviorally react to objects that stimulate less than one-third of the acceptance angle of an individual retinal ommatidium [27,28]. To confirm that flies are capable of perceiving such small objects projected by our LED display, we used a systems identification method [29] to measure the behavioral impulse response generated by the displacement of a single pixel (3.75°). We found robust steering responses to either a single pixel object or two-pixel object displaced in single-pixel increments (Figure 1I inset). Using a display composed of the same LED panels with slightly larger circumference we found that a moving 2.2° object drove LC11 responses well above noise and greater than half-amplitude response to a 4.4° object (Figure 1I), which itself also falls beneath the inter-ommatidial separation angle of 5° [27]. Although similar observations have been documented in hoverflies [30] and dragonflies [31], to our knowledge, this is the first example of simple “hyperacuity” in *Drosophila.* The spatial resolution of LC11 or other similar neurons in the fruit fly lobula could therefore supply behavioral responses to moving objects that are smaller than the acceptance angle of a single ommatidium [28].

We next compared the calcium response dynamics within dendrites, cell bodies, and axon terminals in the same preparation. Comparing the normalized response trajectories demonstrated that dendrite and terminal responses were tightly synchronized (Figure 1J). By contrast, cell bodies exhibited a significantly delayed response onset (Figure 1K) followed by a sustained decay to baseline. These differences may be attributed to the unipolar morphology of invertebrate neurons, in which the cell body is electronically insulated by a long neurite and does not participate in synaptic integration.

Despite the slow response kinetics, imaging from the cell bodies is required to access individual neurons within the palisade of LC11 cells. To map the receptive field of individual LC11s we swept an 8.8° square object in horizontal and vertical directions at each elevation and azimuthal angle, respectively (Figure 2A). For each LC11 recording, peak ΔF/F values were fit to Gaussian functions of azimuth and elevation and used to estimate the two-dimensional spatial receptive field (Figure 2B and S2). We enclosed the spatial receptive field with a contour representing the full-width at 25% max of the Gaussian fits (Figure 2B, see Supplemental Experimental Procedures). We were generally able to record several distinct LC11 cell bodies from each preparation (Figure 2C). Some receptive fields we sampled overlapped, providing functional evidence that individual LC11s have overlapping dendrites (Figure 2C). The average receptive field size was 26.2° by 20.7° with standard deviation of 4.4° and 5.1° respectively (Figure 2D). The functional receptive field size is well below the anatomical dendritic field size (70°-75° by 30°-40°, Figure 1), a property not shared by previously identified neurons in the lamina, medulla or lobula plate [13,32,33].

**Figure 2.**
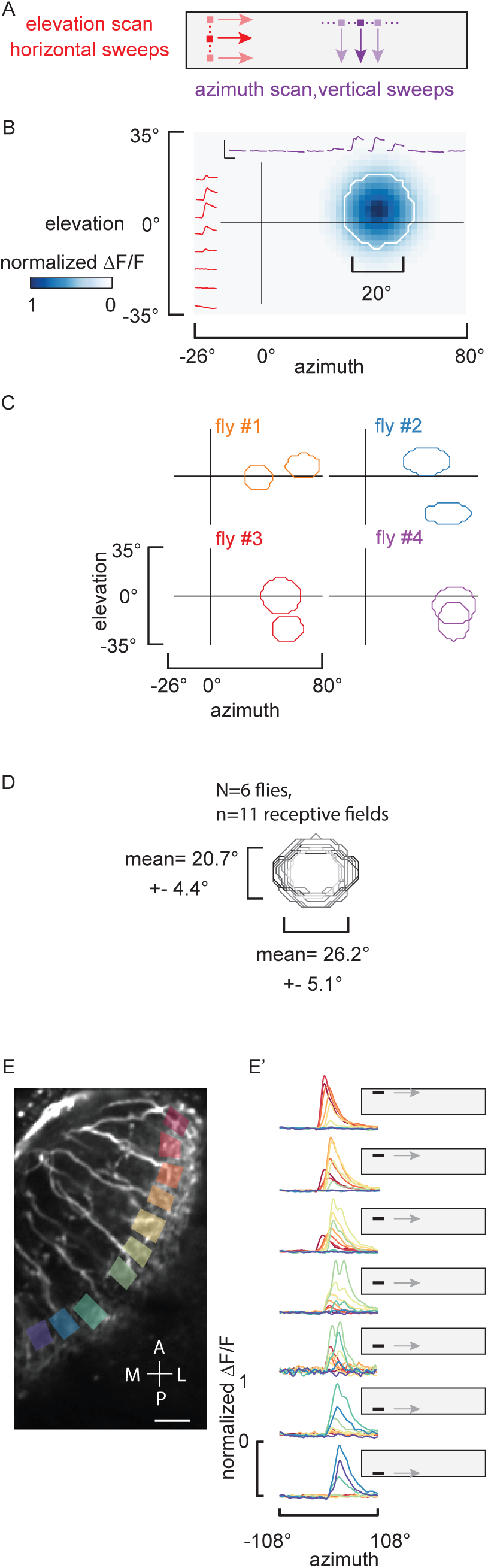
LC11 receptive fields revealed by cell body imaging. A) Schematic of the experimental stimuli used to map individual LC11 receptive fields. An 8.8° dark object moved along non-overlapping trajectories on both horizontal and vertical dimensions at 33 °/sec. B) Reconstructed receptive field of a single LC11. Individual imaging responses from a single LC11 to the horizontal and vertical sweeps shown in red and purple respectively (scale bar represents ΔF/F; y-axis = 0 – 200% and x-axis = 0 – 5 seconds). Reconstructed estimate of a single LC11 receptive field (see Supplemental Experimental Procedures) shown in blue, the full-width at 25% max contour was drawn around the receptive field in white. C) Representative receptive field 25% max contours from different preparations are mapped onto the projection of the visual field. D) 11 receptive fields from 6 flies are overlaid. E) ROIs from separate dendritic compartments are indicated with a different color box. ΔF/F responses to an object swept horizontally at the elevation indicated plotted for all 10 ROIs and overlaid. E’) To facilitate comparison, the responses are normalized to the maximum ΔF/F calcium signal at each ROI. Note that anterior (red) ROIs are activated by object motion across the top of the display, whereas posterior (blue) ROIs are activated by object motion across the bottom of the display. Scale bar for the two-photon image represents 10 um. Abbreviations for anatomical directions; A: Anterior, P: Posterior, M: Medial and L: Lateral.

To assess whether LC11 dendrites sample the full visual field in a retinotopic manner, we imaged from dendritic arborizations of neighboring LC11s simultaneously and presented horizontal object motion at varying elevation angles (Figure 2E). Objects that are presented at higher elevations elicited responses from anteriorly located dendrites (Figure 2E’). As the object moved from higher to lower elevations on the display, the dendritic responses shifted from the dendrites of anterior to posterior LC11 (Figure 2E’) confirming the retinotopic organization of LC11 receptive fields. Given the sluggish response kinetics observed in the cell bodies (Figure 1J), the tight temporal coupling between GCaMP responses in the dendrites and axon terminals (Figure 1J), and the full innervation by individual LC11 cells throughout the output glomerulus (Figure 1B i-ii), we carried out further detailed characterization of LC11 responses from the axon terminals in the VLPR.

The putative neuronal homologues of LC11 in the dragonfly [34] and hoverfly [30] are called small target motion detectors (STMDs) and show a preference for objects darker than the background. To test whether LC11 exhibits a contrast polarity preference, we presented a light object moving across a dark background (ON) and dark object moving on a light background (OFF) (Figure 3A). Our results indicate that LC11 responds to both OFF and ON stimuli yet shows significant preference for OFF stimuli (Figure 3A’, paired t-test p<0.001). Another characteristic feature of STMDs is flicker insensitivity [35]. Stimulating with a stationary object localized within the receptive field that either stepped up in brightness (ON) or stepped down in brightness (OFF) failed to elicit a response in the terminals of LC11 (Figure 3A’’). To test whether the response amplitude has a linear relationship to object contrast polarity we systematically varied the object brightness on a fixed intensity background (Figure 3B). Interestingly, reduction of the OFF-object contrast from 100% to 30% nearly doubled the amplitude of the calcium response (Figure 3B). LC11 responds only weakly to moving ON objects and does not show a significant change in amplitude across contrast (Figure 3B’), suggesting that effect of contrast is OFF object-specific. In flies ON increments and OFF decrements in static light intensity are separated in the lamina and relayed to deeper neuropils via parallel pathways [13,16,19,36]. Layers 2-4 of lobula are innervated by ON-selective Tm3 and OFF-selective Tm4 neurons [37]. Correlating a delayed OFF signal with an un-delayed ON signal arising from a single photoreceptor, i.e. within the same column, fully captured the contrast selectivity of the dragonfly STMD [34].

**Figure 3.**
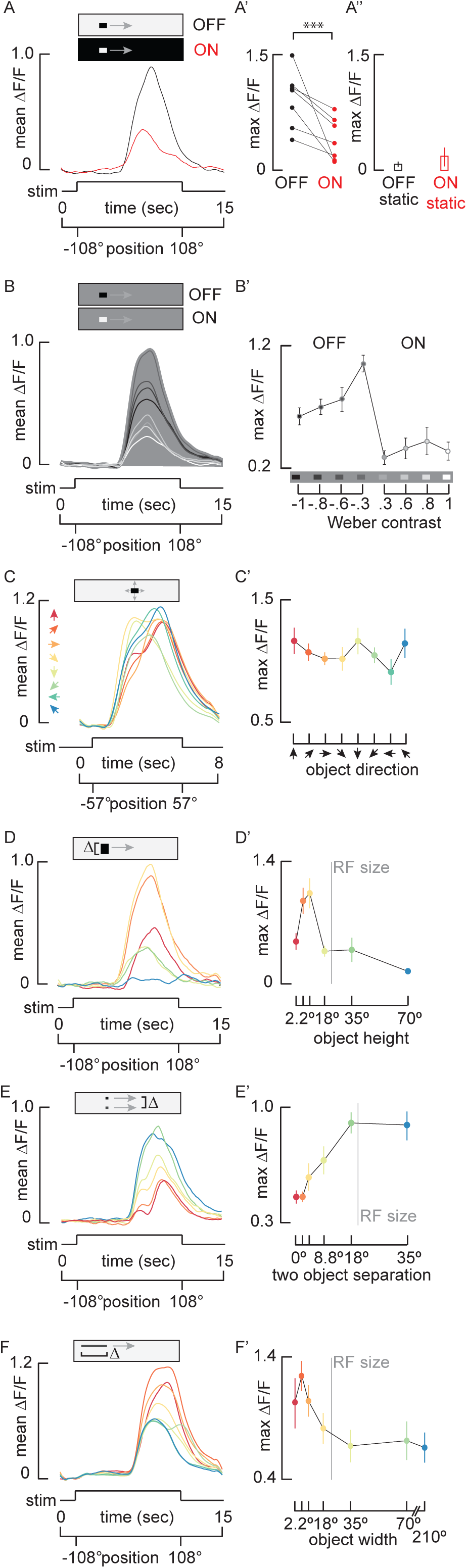
LC11 shows contrast selective object tuning. A) Mean GCaMP6m signal from the LC11 glomerulus in response to a 30°by 8.8° moving ON object (red) and moving OFF object (black, n = 7 flies). A’). Pairwise comparison of maximum ΔF/F of responses from each preparation to both an ON object and an OFF object (***p<0.001, paired t-test, n = 7 flies). A’’)Average of maximum responses (±S.E.M.) of LC11 glomerulus to a stationary 30°by 8.8°OFF and ON object placed within the hotspot of the receptive field (n = 6 flies). B) Mean GCaMP6m signal from the LC11 glomerulus in response to varying contrast objects. Grayscale of the filled area indicates the intensity of the visual background and grayscale of the response line indicates intensity of the stimulus object (n = 9 flies). This display scheme highlights that the most contrasting objects do not elicit the maximum responses from LC11. B’) Average of maximum responses (±S.E.M.) of the LC11 glomerulus to objects of varying contrast. Schematic on the x-axis shows the intensity of the background compared to each object. Weber contrast values are indicated numerically (see Supplemental Experimental Procedures, one-way ANOVA, n = 9 flies). C-F) Average GCaMP6m signals from the LC11 glomerulus to stimulus parameters as indicated. The color map to the stimulus parameter is indicated in C’-F’. C’-F’) Average maximum GcAMP6m (±S.E.M) from the LC11 glomerulus generated from the data in Figure 3C-F, C and C’) LC11 is non-directional. An 8.8° square object was moved in 8 different directions in 45° steps as indicated by color-coded arrowheads (n = 7 flies). D and D’) LC11 is vertically size tuned. A 30° wide object was moved on the same horizontal trajectory, with varied vertical separation: 2.2°, 4.4°, 8.8°, 18°, 35°, 70°, colors mapped to object size in B’ (n = 7 flies). Vertical gray line indicates average receptive field (RF) size (Figure 2). E and E’) LC11 is inhibited by a second object. Two 8.8° square objects moved on parallel trajectories. The distance between them was 0°, 2.2°, 4.4°, 8.8°, 18° and 35°, colors mapped to separation distance in C’ (n = 6 flies). Vertical gray line indicates average RF size (Figure 2). F and F’) LC11 is horizontally size tuned. An 8.8° high object was moved horizontally with varied angular width: 2.2°, 4.4°, 8.8°, 18°, 35°, 70°, 210°, colors mapped to object width in D’ (n = 15 flies). The leading edge of each object appeared on the LED display at the same time. Vertical gray line indicates average RF size (Figure 2).

Given that LC11 is selective for motion rather than flicker (Figure 3A’’), we tested for directional tuning by moving an 8.8° by 8.8° object in eight different directions. LC11 is not significantly selective for motion direction (Figure 3C and C’, one-way ANOVA n.s.), although shows a slight trend for vertical directions, which is consistent with a previous report that measured membrane potential at the cell body and showed subtle preference for downward motion of a large object [38]. A similarly sized object elicits only a weak increase in GCaMP6 fluorescence (Figure 1H).

The selectivity for small moving objects over elongated bars suggests that LC11 is size tuned. The classical mechanism for size tuning is end-stopped inhibition, a hypercomplex property in which an elongated contour stimulates the inhibitory end zones of a receptive field with an excitatory center [1]. We parameterized the vertical dimension of a horizontally moving object of fixed width. The optimum LC11 response occurs for a vertical extent of roughly 9°, which is 1/4^th^ the vertical projection of the anatomical receptive field or ½ the functional receptive field (Figure 3D). The response magnitude asymptotes as the vertical size spans one functional receptive field (Figure 3D’).

The diminished response observed in LC11 for an elongated object has also been observed in dragonfly STMDs [39]. Owing to the close match between object size tuning and the functional receptive field, we reasoned that the end-stopped property of LC11 could be shaped by lateral inhibition generated by nearest neighbor LC11s. If so, then the end-stopped inhibition generated by two nearby objects should be fully relieved once they are separated by one receptive field increment. By presenting two 8.8° square objects moving horizontally, we confirmed that response amplitude increased with increasing object separation (Figure 3E). The separation distance had no effect on response amplitude once the two objects were separated by ~18°, roughly the size of a single LC11’s functional receptive field (Figure 3E’). Furthermore, separation distances near the preferred object size (9°) led to the biggest modulation in response amplitude. These results support the hypothesis that end-stopped inhibition could originate from nearest neighbor LC11s.

In addition to classical end-stopped properties, we also observed a peculiar increase in response amplitude when the contrast of a 30° by 8.8° object was systematically decreased (Figure 3B’). Notably, the object size in these experiments (30°) is larger than the estimated receptive field size (26°). It is likely that a 30° object traverses both excitatory receptive field and inhibitory end zones. It has been proposed previously that end stopping only works when the edges of the object have the same contrast as the center [40]. Reduced contrast of a sufficiently large object may therefore release the inhibition generated by the object edges, resulting in increased response amplitude observed in LC11 at reduced contrast (Figure 3B’).

One consistent characteristic of the response to two objects is a ‘double peak’ for object separation less than 18° (Figure 3E). We did not observe the bi-phasic response dynamics when presenting the same stimuli in the reverse direction, during which the targets swept from the back-to-front direction (Figure S3). Thus, we attribute the ‘double peak’ phenomenon either to subtle differences in the spatial distribution of LC11 receptive fields converging in the axon terminals, or to differences in the spatial properties of inhibition between the lateral-ventral field of view by comparison to the frontal-dorsal field.

To test whether LC11 is size-tuned in the dimension parallel with the axis of motion, we presented objects of fixed vertical height moving horizontally with varying horizontal widths. As the width of the object increased, the response amplitude peaked near 4.4° (Figure 3F). The peak amplitude decreased until the width of the object was approximately the equivalent of one LC11 receptive (~26°) (Figure 3F’). Note that the end-stopped inhibition generated by an elongated object oriented perpendicular to its axis of motion (Figure 3D) is stronger than the inhibition generated by an object oriented parallel to its axis of motion (Figure 3F). This would be expected because the former activates more LC receptive fields, and thereby generates more spatial inhibition, than the latter.

For the experiment in which we parameterized the vertical size of the horizontally moving object (Figure 3D), we would expect each object to sweep through different numbers of LC11 receptive fields at different times and thereby differences in the onset timing of the GCaMP signals in the glomerulus (Figure 3D). By contrast, for the horizontally moving object, the spatial extent of the stimulus orthogonal to the motion vector is invariant and therefore stimulates the same ensemble of LC11 receptive fields for each stimulus. Hence, the onset delay is invariant for this experiment (Figure 3F).

Taken together, the results presented thus far implicate end-stop lateral inhibition in sculpting object response properties, suggesting that LC11 may receive GABAergic input. Previous immunohistochemical studies of the lobula indicate that antibodies targeting GABAergic signaling pathways such as glutamic acid decarboxylase (GAD), the GABA_A_ receptor subunit RDL, and GABA itself, are present in a layer-specific manner [41,42]. In particular, an enrichment of GABAergic neurotransmission, indicated by dense labeling of all three antibodies, is observed in layers 2 and 3 of the lobula. Our own co-labeling experiments indicate a strong overlap between the dendritic arborizations of LC11 and enriched vesicular GABA transporter (VGAT) staining (Figure 4A and C). In contrast, the dendrites of LC11 are spatially-excluded from layer 1, known to be enriched with cholinergic signaling within T5 cells (Figure 4B and D) [43].

**Figure 4.**
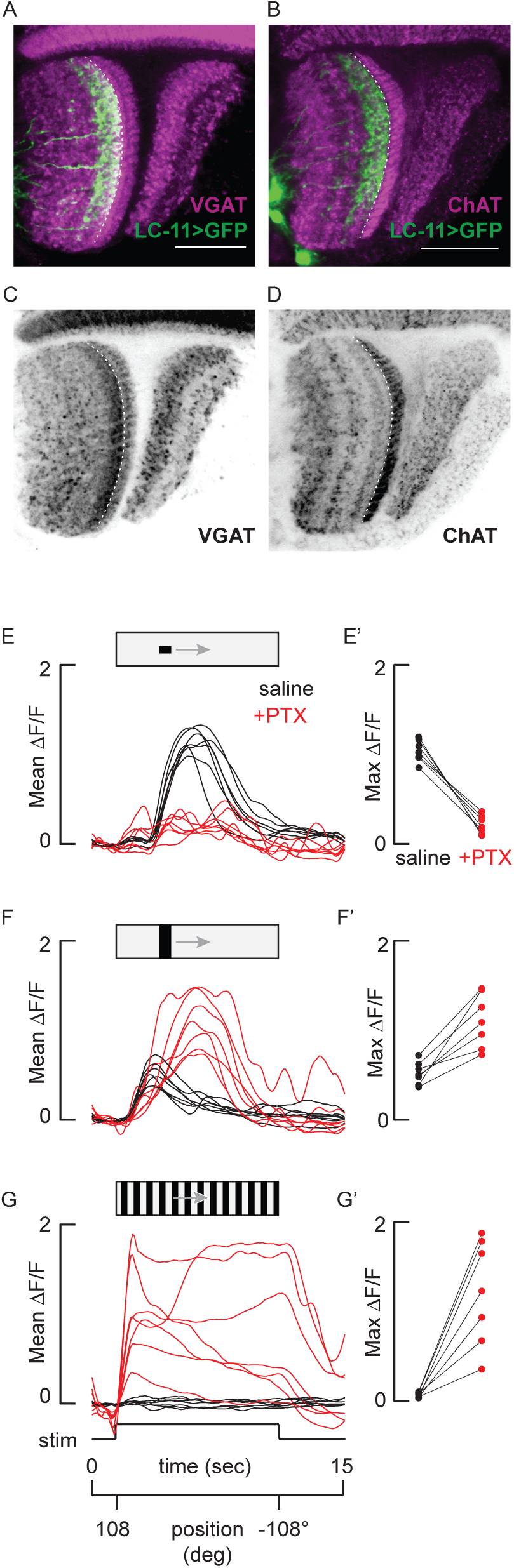
Object selectivity in LC11 is driven by synaptic inhibition. A and B) LC11 dendritic layer is enriched with vesicular GABA transport (VGAT), and the adjacent presumably presynaptic layer with choline acetyltransferase (ChAT). Dorsal view of GFP labeled LC11 neurons (green) co-labeled with either anti-ChAT (A, magenta) or VGAT (B, magenta). Dashed line indicates the border between the first and second lobula strata. Scale bars 25 um. C and D) Layering of VGAT and ChAT are highlighted. Same labeling as in A and B, without LC11 overlaid. Scale bars are 25 um. E – G) Inhibition sculpts object responses and inhibits bar and grating responses. Average DF/F glomerular responses of LC11 from n = 7 flies in response to a 30°by 8.8° object (E), a 30° by 70° bar (F) and a wide-field grating (G) with (red) or without (black) 10 um picrotoxin (10 uM). E’ – G’) Comparison of maximum ΔF/F of values from each fly (E - G) with (red) or without (black) picrotoxin.

To test for functional consequences of GABAergic inhibition on the response properties of LC11, we imaged from the LC11 glomerular outputs before and after blocking GABA_A_-mediated inhibitory currents with picrotoxin (PTX). Prior to PTX application, LC11 responded robustly to a small object, showed a slight excitation in response to the onset of the bar, and was not at all excited by a wide-field grating (Figure 4E,F and G). Applying PTX in the perfusion saline within the same recording preparations resulted in strongly reduced object responses (Figure 4E), large amplitude bar responses (Figure 4F), and large amplitude sustained responses to wide-field motion (Figure 4G). Remarkably, PTX-dependent loss of not only selectivity, but also sensitivity to a small object was measured in every recording preparation (Figure 4E’, F’ and G’). Thus, inhibitory currents not only ‘end-stop’ LC11 for size tuning, these currents also actively sculpt the detection of small objects. By contrast, figure detecting cells (FD) of the lobula plate in larger flies are excited by small-field gratings and receive GABAergic wide-field inhibition, yet FD continues to respond to small-field motion under GABA-blockade [44]. In STMDs, small object tuning has been attributed to lateral inhibition at the level of pre-synaptic neighboring ON-OFF channels that are correlated and summed [34,45]. However, in the absence of inhibition the model predicts that STMDs would be driven by dark edges of any size. Our PTX results are the first to demonstrate that inhibition is required for small object detection itself (Figure 4E).

Taken together with the findings fact that LC11 spans 14-15 columns of the visual world yet has a functional receptive field roughly 4-6 columns and peak size tuning less than a single column, suggests complex inhibitory spatial interactions. We propose that an excitatory-center inhibitory-surround mechanism, driven by nearest-neighbor inhibition, spatially sharpens the receptive field of LC11 making it both sensitive and selective for small contrasting objects.

Although we provide ample amount of physiological data to illustrate that LC11 is an object detector we have not been able to pinpoint the contribution of this cell type to behavior. We tested for LC11 contribution to small object avoidance in flight [46], but did not observe a clear phenotype (data not shown). LC11 could instead contribute to different small-object behaviors such as courtship [11,47]. Alternatively, the complex spatial receptive field interactions of LC11, activating or inactivating the entire population may produce spurious results that avoid detection by our assay. Finally, LC11 is one of over 16 types of VPNs that innervate similar input layers in the lobula [5,21], and other optic glomeruli are also likely to be small object detectors. Nevertheless, we show extensive physiological characterization and mechanistic analysis of the object selectivity of a visual projection neuron in *Drosophila*, which shares functional properties of object detectors in other insects and vertebrates.

## Author Contributions

M.F.K. and M.A.F. designed the research and wrote the manuscript. M.F.K. conducted the experiments and analyzed the data.

## Acknowledgements

We thank Jaison Omoto for carefully reading the manuscript, Jacob Aptekar for experimental help and contribution to subpanel of Figure 1I, Volker Hartenstein for confocal microscope use and Frye lab members for helpful discussion, Aljoscha Nern and Michael Reiser for lobula layer identification and kindly discussing unpublished results, David Krantz for the VGAT antibody. This research was supported by the UCLA Edith Hyde Fellowship (M.F.K.), and National Institutes of Health EY026031 (M.A.F.).

